# Optimizing clinical interpretability of functional evidence in epilepsy-related ion channel variants

**DOI:** 10.1101/2024.05.09.593343

**Authors:** Shridhar Parthasarathy, Stacey R. Cohen, Eryn Fitch, Priya Vaidiswaran, Sarah Ruggiero, Laina Lusk, Victoria Chisari, David Lewis-Smith, Stephan Lauxmann, Christian M. Boßelmann, Christopher H. Thompson, Eric R. Wengert, Ulrike Hedrich, Shiva Ganesan, Ganna Balagura, Roland Krause, Julie Xian, Peter Galer, Manuela Pendziwiat, Eduardo Perez-Palma, Mauno Vihinen, Jennifer Hart, Melissa J. Landrum, Dennis Lal, Edward C. Cooper, Holger Lerche, Ethan Goldberg, Andreas Brunklaus, Carlos G. Vanoye, Stephanie Schorge, Alfred L. George, Ingo Helbig

**Affiliations:** Division of Neurology, Children’s Hospital of Philadelphia, Philadelphia, PA, 19104, USA; The Epilepsy NeuroGenetics Initiative (ENGIN), Children’s Hospital of Philadelphia, Philadelphia, PA, 19104, USA; Department of Biomedical and Health Informatics (DBHi), Children’s Hospital of Philadelphia, Philadelphia, PA, 19146, USA; Medical Scientist Training Program, Perelman School of Medicine, University of Pennsylvania, Philadelphia, PA, 19146, USA; Temple University, Lewis Katz School of Medicine, Philadelphia, PA, 19140, USA; FutureNeuro SFI Research Centre, RCSI University of Medicine and Health Sciences, 123 St Stephen’s Green, Dublin 2, Ireland; Translational and Clinical Research Institute, Newcastle University, Newcastle-upon-Tyne NE2 4HH, UK; Department of Neurology and Epileptology, Hertie Institute for Clinical Brain Research, University of Tübingen, Tübingen, Germany; Department of Pharmacology, Northwestern University Feinberg School of Medicine, Chicago, IL, 60611, USA; Department of Neurosciences, Rehabilitation, Ophthalmology, Genetics and Maternal and Child Health, University of Genoa, Genoa, Italy; Child Neurology and Psychiatry, Istituto G. Gaslini, Genoa, Italy. Member of ERN EpiCARE.; Luxembourg Centre for Systems Biomedicine, Université du Luxembourg, Esch-sur-Alzette, Luxembourg; Medical Scientist Training Program, Johns Hopkins School of Medicine, Baltimore, Maryland, 21205, USA; Department of Bioengineering, School of Engineering and Applied Sciences, University of Pennsylvania, Philadelphia, Pennsylvania, 19104, USA; Center for Neuroengineering and Therapeutics, University of Pennsylvania, Philadelphia, Pennsylvania, 19104, USA; Institute of Clinical Molecular Biology, Christian-Albrechts-University of Kiel, Kiel, Germany; Universidad del Desarrollo, Centro de Genética y Genómica, Facultad de Medicina Clínica Alemana, Santiago, Chile; Department of Experimental Medical Science, Lund University, Lund, Sweden; National Center for Biotechnology Information, National Library of Medicine, National Institutes of Health, Bethesda, Maryland, 20894, USA; Epilepsy Center, Neurological Institute, Cleveland Clinic, Cleveland, 44195, USA; Stanley Center for Psychiatric Genetics, Broad Institute of MIT and Harvard, Cambridge, Massachusetts, 02142, USA; Departments of Neurology, Neuroscience, Molecular and Human Genetics, Baylor College of Medicine, Houston, Texas, 77030, USA; Department of Neurology, University of Pennsylvania Perelman School of Medicine, Philadelphia, PA, 19104, USA; The Paediatric Neurosciences Research Group, Royal Hospital for Children, Glasgow, UK; Institute of Health and Wellbeing, University of Glasgow, UK; Department of Pharmacology, University College London School of Pharmacy, London, UK

## Abstract

Variants in genes encoding the voltage-gated ion channels are among the most common monogenic causes of epilepsy and neurodevelopmental disorders. Functional effects of a variant are increasingly important for diagnosis and therapeutic decisions. To incorporate knowledge regarding functional consequences in formal clinical variant interpretation, we developed an approach for evaluating multiple functional measurements within the Bayesian framework of the modified ACMG/AMP guidelines. We analyzed 216 functional assessments of 191 variants in *SCN1A* (n=74), *SCN2A* (n=66), *SCN3A* (n=18), and *SCN8A* (n=33). Of 20 commonly measured biophysical parameters, the most frequent drivers of overall functional consequence were persistent current (f=0.54), voltage dependence of activation (f=0.51), and voltage dependence of fast inactivation (f=0.40) for gain-of-function and peak current (f=0.87) for loss-of-function. By comparing measurements of 23 benign variants, we determined thresholds by which published data on these four parameters confer *Strong* evidence of variant pathogenicity (likelihood ratio > 18.7) under the ACMG/AMP rubric. Similarly, we delineated evidence weights for the most common epilepsy-related potassium channel gene, *KCNQ2*, through reports of 80 pathogenic and 24 benign variants, accounting for heterozygous and homozygous experimental conditions. We collected the resulting categorization of functional data into FENICS, a biomedical ontology of 152 standardized terms for coherent annotation of electrophysiological results. Across 271 variants in *SCN1A/2A/3A/8A* and *KCNQ2*, 1,731 annotations are available in ClinVar, facilitating use of this evidence in variant classification. In summary, we introduce and apply an ACMG/AMP-calibrated framework for electrophysiological studies in epilepsy-related channelopathies to delineate the impact of functional evidence on clinical variant interpretation.

## Introduction

Variation in ion channel genes is the most common cause of human epilepsy and neurodevelopmental disorders.^1,2^ Individuals with disease-causing variants in *SCN1A*, *SCN2A*, *SCN3A*, and *SCN8A*, genes encoding the neuronally expressed voltage-gated sodium channels (Na_V_), can have one of many, often severe, clinical presentations, including Dravet syndrome, genetic epilepsy with febrile seizures plus (GEFS+), brain malformations, autism spectrum disorder, and early-onset epileptic encephalopathy.^3-7^

Across these genetic disorders, determining a molecular diagnosis depends upon standardized variant interpretation. The most widely used framework for classifying sequence variants is the American College of Medical Genetics and Genomics and Association for Molecular Pathology (ACMG/AMP) criteria, which were initially published in 2015.^8^ The Clinical Genome consortium (ClinGen) has since clarified these rules using a systematic, Bayesian approach that provides quantitative thresholds for strength of evidence and overall variant classification.^9^ In parallel with the development of these criteria, the number of variants of uncertain significance (VUS) has grown exponentially as clinical genetic testing rapidly expands;^10,11^ efforts to resolve variant pathogenicity are therefore increasingly critical.^12^

Functional studies that provide insight into the underlying biology are a key step in linking genotype to disease. Within the ACMG/AMP rubric, a sufficiently validated clinical functional assay can contribute evidence of pathogenicity at a *Strong* level, comparable to the criterion for a verified *de novo* variant.^8,9^ The ClinGen modifications to this PS3 criterion provide for more rigorous quantification, stratification of functional evidence across multiple levels of strength, and avenues for inclusion of experiments performed on a research basis.^13^ In the voltage-gated channelopathies, this has already permitted detailed evaluation of a functional assay for *KCNH2*, a gene linked to long QT syndrome.^14^ However, this represents only one of numerous disease-linked channels, and there is currently limited scalability in developing new calibrations of functional study data from scratch.

As an alternative resource to new assessments of functional assays, there is a wealth of existing electrophysiological research data reported from research settings. Voltage clamp experiments are the standard in research for the functional study of voltage-gated ion channel variants, and emerging technologies are rapidly increasing scale and throughput of these assays.^15-18^ Rigorously leveraging these data within the ACMG/AMP framework would increase available evidence for variant interpretation. However, reporting of results is sparse and heterogeneous. Electrophysiological studies can be performed in a variety of model systems with multiple methods of measurement and a wide range of output formats.^16,19,20^ Moreover, investigators may not study the same biophysical parameters for all variants. A wide range of effects can be included in a study, such as different quantifications of current density and gating, some of which are not commonly measured.

Beyond diagnosis and variant classification, electrophysiology can be used to elucidate clinical trajectories and guide therapeutic decision-making. For example, overall functional consequences are increasingly correlated with emerging clinical stratifications, such as the gain-of-function spectrum in *SCN1A*.^21,22^ We and others also have demonstrated a strong link between electrophysiological findings and the phenotypic landscape of *SCN2A*-related disorders.^6,23^ Yet, the bulk of this prior work only distinguishes high-level variant descriptions—namely, overall gain-of-function, loss-of-function, mixed/uncertain, or normal—partly as a consequence of the diversity in reporting of functional data. No framework exists to adequately capture the more granular properties measured in these studies at a larger scale, which would allow for mechanistic insights that inform precision therapy approaches.

Here, we aim to address these challenges by introducing the Functional Electrophysiology Nomenclature for Ion Channels (FENICS), a biomedical ontology of functional changes to voltage-gated ion channels, to standardize use of electrophysiological experiments in clinical variant interpretation. We curated 216 published experiments in *SCN1A/2A/3A/8A* with 1,484 functional annotations and established thresholds that are aligned with the ClinGen modified ACMG/AMP criteria across the commonly measured electrophysiological parameters. By extending this framework to the potassium channel *KCNQ2*, we show that our approach can be applied broadly across channelopathies towards guiding diagnosis, and eventually, future precision medicine avenues.

## Methods

### Collection of ion channel variant functional assessments

We reviewed the literature published until October 2023 for electrophysiological assessments of the genes encoding epilepsy-related voltage-gated sodium channel pore-forming subunits, namely *SCN1A*, *SCN2A*, *SCN3A*, and *SCN8A*. We defined a single assessment as the set of all measurements of a single variant reported within a single publication. We retrieved 216 total functional assessments in the literature for 191 missense variants in *SCN1A* (n=74), *SCN2A* (n=66), *SCN3A* (n=18), and *SCN8A* (n=33, **Figure 1, Table S1**). We included all experimental data in variants with multiple studies, distinguishing the individual assessments. Although we curated experiments performed on the fetal isoform of *SCN2A*, we excluded these results from subsequent analysis, but we retained the parallel studies performed on the adult isoform. We also documented methodological information from each experiment, such as the cell lines used. Additionally, we obtained data on *KCNQ2* from a recent study of clinical and population variants using automated patch recording.^16^ In that study, all recordings were made using a cell line expressing an electrically silent *KCNQ3*/Kv7.3 subunit, and *KCNQ2* variants were introduced by transient transfection, either alone (“homozygous” expression) or in a 1:1 ratio with wild-type (WT) *KCNQ2* to mimic the heterozygous state.

**Figure 1.**
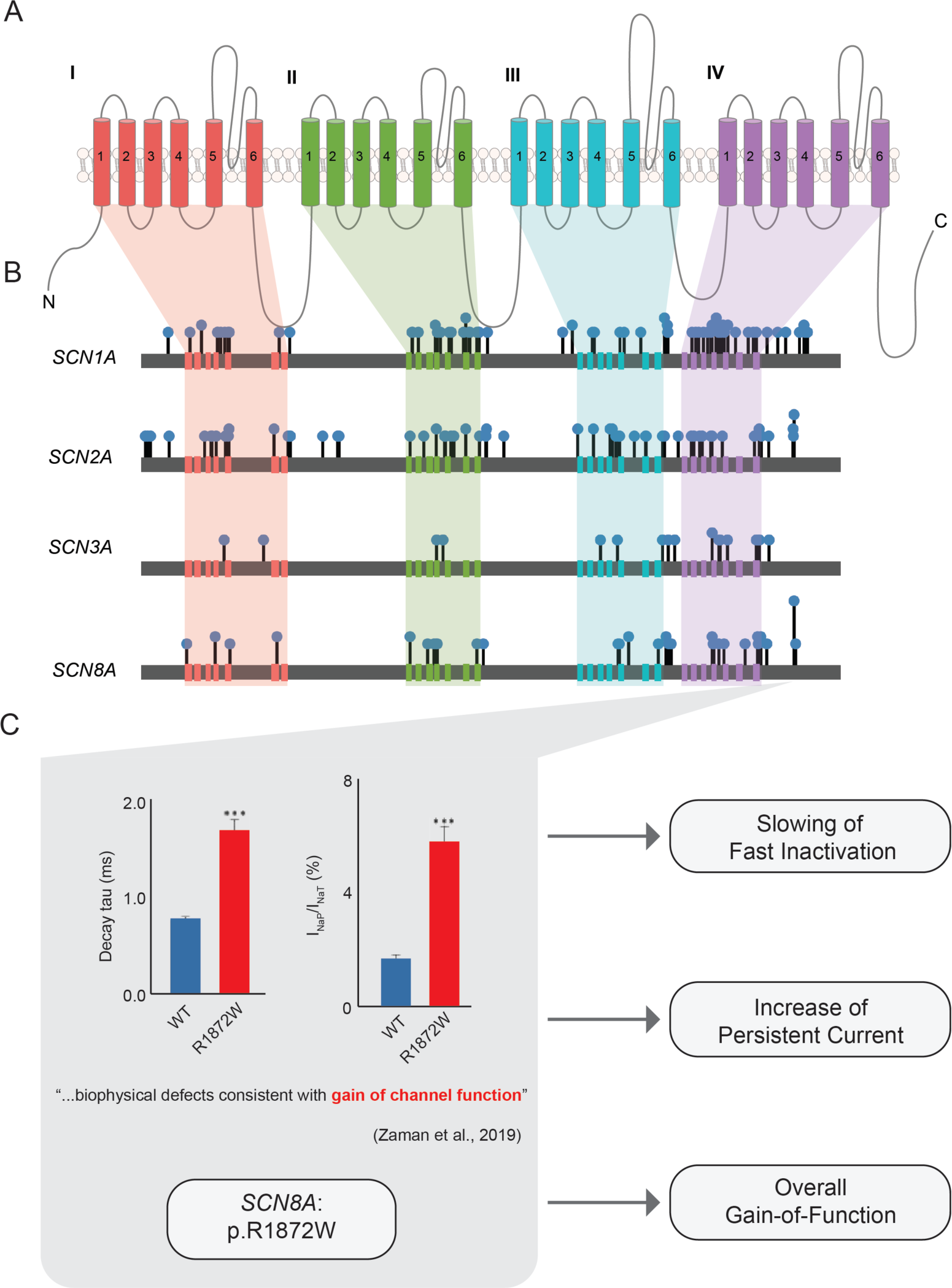
Review of 216 experiments in 191 *SCN1A/2A/3A/8A* variants. (A) The common structure of the voltage-gated sodium channel, highlighting four homologous domains, each with six transmembrane segments including the S4 voltage sensor and S5-6 pore loop. (B) Map of functionally studied variants in the sodium channels, showing number of distinct functional studies (height) and overall functional effect (color). For 168/191 variants, only a single experimental assessment has been performed. The most analyzed variant position is *SCN8A*:p.R1872Q/W/L, for which 7 independent experiments have been performed. (C) Example categorical mapping of functional data on *SCN8A*:p.R1872W.34 Where available, changes to each of 22 biophysical parameters were recorded along with the overall effect of the variant.

### Calibration of functional evidence for the ACMG/AMP PS3 criterion

We developed a framework in line with the revised ClinGen guidelines for assessing levels of strength of evidence for variant pathogenicity. In particular, for sodium channels, we chose benign and pathogenic control variants according to the specifications of the ClinGen Epilepsy Sodium Channel Variant Curation Expert Panel (VCEP) (accessed via <u>cspec.genome.network/cspec/ui/svi/affiliation/50105</u>). Twenty-three benign variants in *SCN1A* were studied by voltage clamp electrophysiology and designated benign controls (unpublished data). These variants met the BA1 population frequency threshold criterion as per their allele frequencies in gnomAD v2.1.1.^24^ Sixty-three variants that were likely pathogenic or pathogenic independent of functional testing had available electrophysiological data and were therefore considered pathogenic controls. We determined threshold values for each of four parameters: peak current density, voltage dependence of activation, voltage dependence of fast inactivation, and persistent current. These parameters were chosen as they represent the most common changes identified in functionally altered disease-causing epilepsy sodium channel variants (**Figure 2**). Peak current density and persistent current were quantified as a ratio relative to WT controls from the same study. Voltage dependence of activation and fast inactivation were documented as shifts in mV, and thresholds were based on the absolute value of deviation from WT. In experiments where parameter values for mutant channels were reported as not different from WT controls, but where raw quantities were not available, we used the value from the corresponding WT controls.

**Figure 2.**
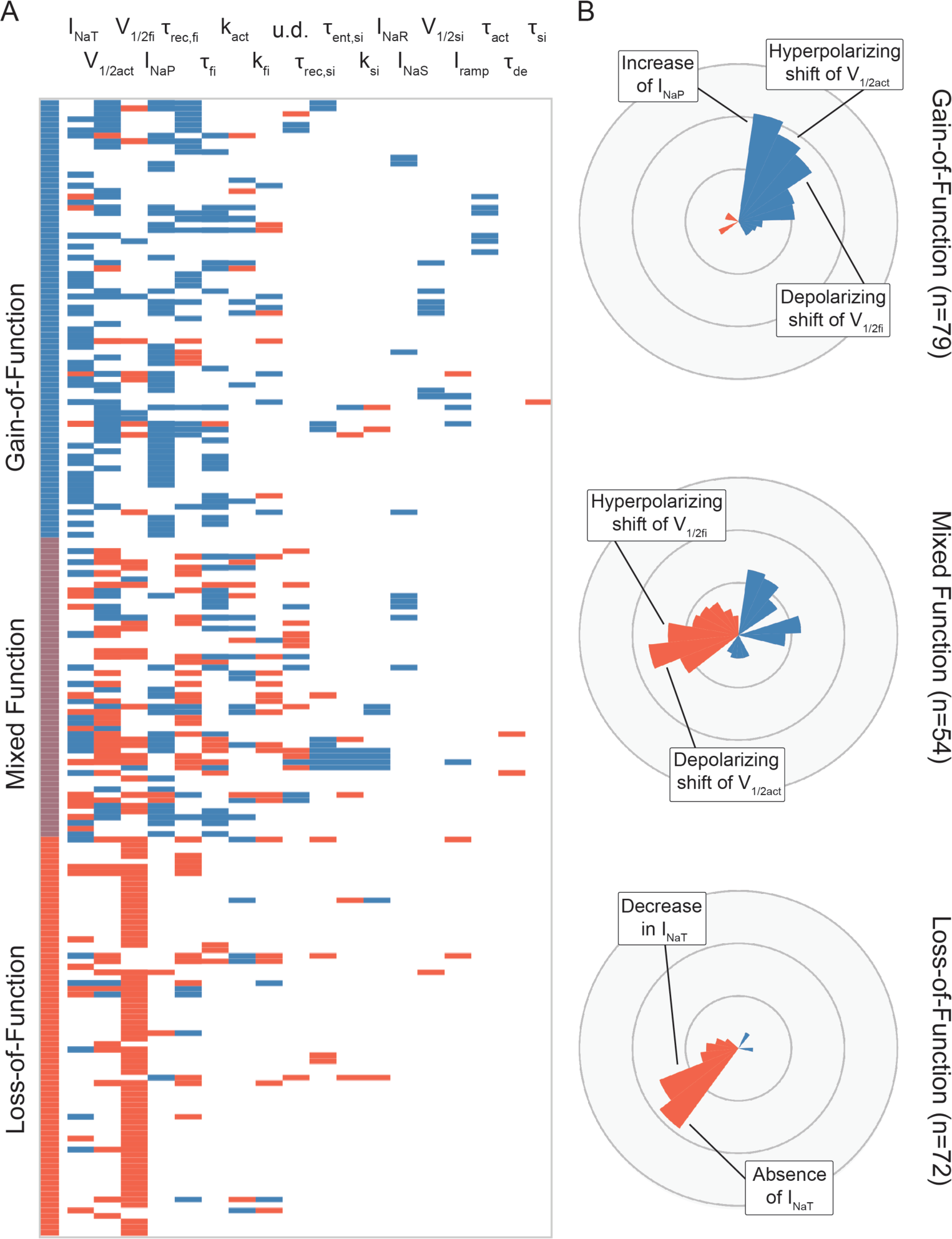
Heterogeneous functional effects across ion channel experiments. (A) Effects on individual biophysical parameters showing gain-of-function (blue), loss-of-function (red), and normal or unmeasured (gray) defects across 19 parameters (rows, ordered by decreasing number of available measurements) and 216 variant assessments, highlighting the complex biophysical landscape of these variants. Variants frequently exhibit properties leading to both gain- and loss-of-function. (B) Distribution of functional effects in overall gain-of-function, loss-of-function, and mixed-function variant assessments. Bars indicate frequency of an effect in the respective subgroup. INaT = peak (transient) current, V1/2act = voltage dependence of activation, V1/2fi = voltage dependence of fast inactivation, INaP = persistent current, υrec,fi = time constant of recovery from fast inactivation, υfi = time constant of fast inactivation, kact = slope factor of activation, kfi = slope factor of fast inactivation, u.d. = decay in current amplitude from use dependence, υrec,si = time constant of recovery from slow inactivation, υent,si = time constant of entry into slow inactivation, ksi = slope factor of slow inactivation, INaR = resurgent current, INaS = subthreshold current, V1/2si = voltage dependence of slow inactivation, Iramp = ramp current, υact = time constant of activation, υde = time constant of deactivation.

As previously described for *in silico* predictors of variant pathogenicity,^25^ we computed a positive local likelihood ratio (LR^+^) for deviation beyond every possible threshold value within our dataset, and where zero false positives were observed, a value of one was used instead. Then, we determined minimum deviations achieving *Strong*, *Moderate*, and *Supporting* evidence levels. However, as each parameter in the voltage clamp assay is not strictly independent, these values are complicated by the presence of incorrect false negatives, i.e., pathogenic variants which are “normal” with respect to one parameter but highly abnormal in at least one other parameter. For example, one report of the *SCN2A* p.R1882Q variant showed a peak current of 100% WT but persistent current of more than 200% WT (**Figure 3A, Table S2**).^26^ It would be inappropriate to classify this variant as “normal” with respect to the overall voltage clamp assessment when evaluating thresholds for peak current. To adjust for this, we iterated the above computation of likelihood ratios for each parameter and determined ACMG/AMP-compatible thresholds for each parameter, filtering out such incorrect false negatives at each step using a conservative approach as follows. After each iteration, the threshold for each parameter achieving the highest likelihood ratio was taken as a filtration cutoff. Variants were excluded from the subsequent iteration of computing likelihood ratios for a given parameter if they were (1) “normal” as per the threshold being tested AND (2) abnormal beyond the filtration cutoff in a different parameter. Likelihood ratios were again computed, and this process was repeated until threshold and likelihood ratio values converged for all parameters for all evidence levels.

**Figure 3.**
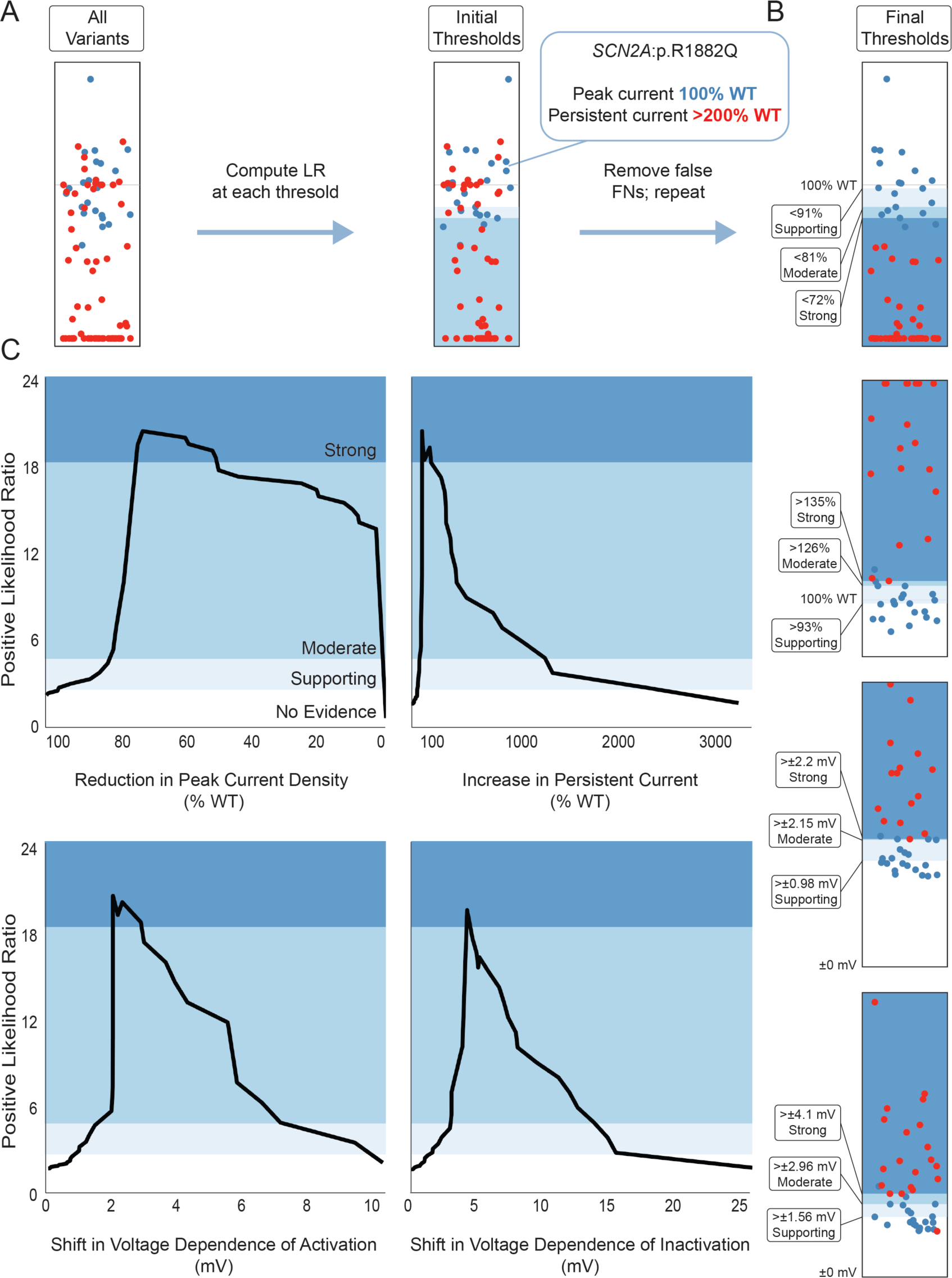
Calibration of voltage clamp measurements *SCN1A/2A/3A/8A* for the modified ACMG/AMP criteria. (A) Example mapping of peak current density. Since the sum total of these parameters represents a single functional assay, likelihood ratios (LR) were computed as previously described, but iteratively for each parameter to remove “false negative” (FN) variants exhibiting other strong biophysical defects. (B) Final ACMG/AMP evidence thresholds for four parameters with sufficient data; dots represent benign (blue, n=23) and pathogenic (red, n=63) controls. (C) Positive likelihood ratios traced across every possible threshold in the final dataset. A *Strong* level of evidence is achievable for each of the parameters analyzed.

We repeated this analysis for variants in *KCNQ2*, choosing control variants as in the sodium channel case. Unlike sodium channels, however, voltage-gated potassium channel pores are assembled as tetrameric complexes., Therefore, pathogenicity of heterozygous disease-causing variants in *KCNQ2* can arise not only from to loss- or gain-of--function, but also through dominant-negative effects.^27-29^ Accordingly, electrophysiological experiments on *KCNQ2* frequently include the paralogous subunit *KCNQ3* and measurements in homozygous and heterozygous conditions in order to capture the range of functional subunit configurations and consequences expected to occur in vivo.^16^ Hence, we calibrated separate sets of severity thresholds for experiments mimicking the heterozygous and homozygous states and for co-expression of *KCNQ2* variants with WT *KCNQ3* and *KCNQ2* subunits. For some variants, certain values were available in the homozygous but not heterozygous state, typically because these homozygous measurements, which are expected to be more extreme than heterozygous, showed no significant difference from WT. For such variants, the homozygous measurements were used as an upper bound, representing the most extreme change that can be expected for a heterozygous experiment on the same variant. We performed evidence threshold computation for *KCNQ2* for peak current density, voltage dependence of activation, and time constant of activation.

### Construction of the FENICS ontology to describe variant experimental results

One challenge in obtaining and using reported voltage clamp data in the clinical genetics setting is the heterogeneous documentation of these results. For example, voltage dependence of fast inactivation may be alternatively referred to with such variable names as “voltage dependence of steady-state inactivation,” “V1/2 inactivation,” or, rarely, “voltage dependence of channel availability.” The same “voltage dependence of channel availability” has, in other contexts, denoted “voltage dependence of activation.” To facilitate accurate use of these data in clinical variant interpretation, we developed a standardized biomedical ontology to which threshold values could be mapped and publicly, centrally documented.

Our ontology consists of a hierarchical framework called the Functional Electrophysiology Nomenclature for Ion Channels (FENICS). We identified commonly measured biophysical parameters and defined categories based on the direction of a difference in each, typically using “increase” versus “decrease” to indicate that a parameter was measured as greater or lesser than WT, respectively. Each of these changes were further subclassified by severity, typically “mild,” “moderate,” and “severe,” with ion channel-specific thresholds established by our ClinGen evidence calibration when applicable, and electrophysiologist expert consensus when not applicable. As a result, this ontology allows for standardized annotation of specific biophysical defects, such as “moderate decrease in peak current,” in a hierarchical manner consistent with existing ontologies such as the Human Phenotype Ontology (HPO) and VariO (**Figure 5A**).^30,31^ We created an independent, parallel branch of the FENICS ontology for use dependence^32^ and ramp current,^33^ as these represent more complex biophysical features of the ion channel that cannot be fully captured by single biophysical properties.

**Figure 4.**
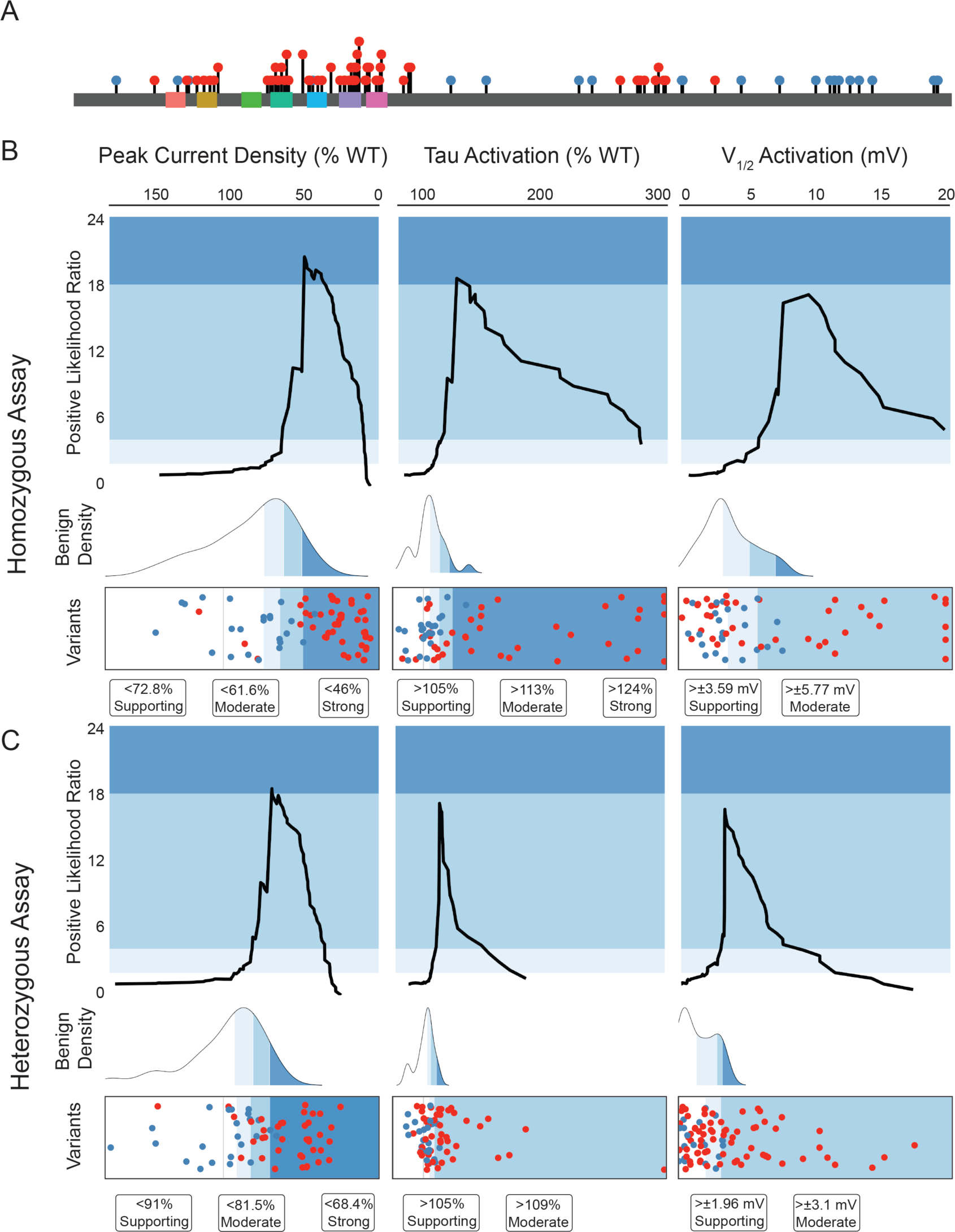
Computation of ACMG/AMP-compatible thresholds for experiments in *KCNQ2*. (A) Mapping of population (blue, n=24) and disease-causing (red, n=80) variants to the KV7.2 channel. Kv7.2 subunits have 6 transmembrane segments, and a long C-terminal intracellular domain. Channels assemble as tetramers including Kv7.2 and Kv7.3 subunits. (B) Likelihood ratios, distribution of benign variant measurements, and final variant evidence thresholds in the homozygous condition for peak current, time constant of activation, and voltage dependence of activation. (C) Likelihood ratios, distribution of benign variant measurements, and final variant evidence thresholds in the heterozygous condition for the same parameters. While voltage dependence of activation in either state and time constant of activation in the heterozygous state do not have *Strong* evidence thresholds, an estimate of a corresponding “severe” categorization can be derived from the 5th percentile of benign variant values.

**Figure 5.**
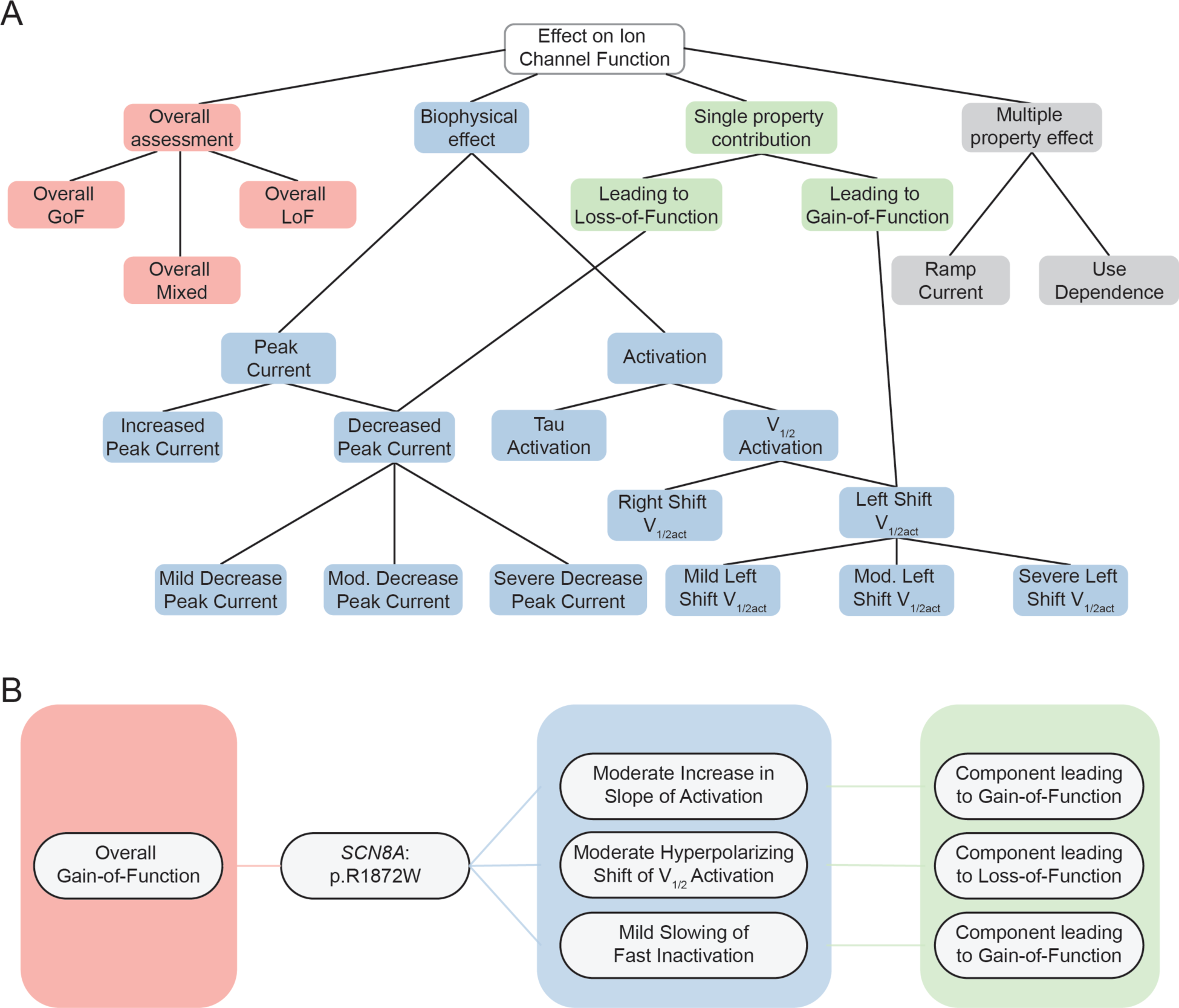
Assembly of expert consensus and ACMG/AMP calibration into the FENICS biomedical ontology. (A) Schematic showing a subset of FENICS. Individual biophysical parameters are subcategorized into directional shifts and ACMG/AMP-compatible levels of severity, which also relate to their contribution to gain- or loss-of-function. Distinct subontologies for more compound parameters and the overall functional consequence also exist. (B) Example translation to FENICS of functional evidence for *SCN8A*:p.R1872W. Annotation of individual parameter changes can automatically map to their functional contribution, and a separate overall effect is also annotated.

For each directional change to a parameter in our dictionary, such as “slowing of fast inactivation,” we assigned a parent term describing its contribution to function, such as “component leading to gain-of-function.” We also included information about overall functional effect of the variant in a separate sub-dictionary to preserve investigators’ overall conclusions, independent of individual parameters (**Figure 5B**).

We translated functional data on selected variants during the development of the dictionary in an iterative manner until the overall dictionary was determined to describe the functional effects sufficiently by an interdisciplinary panel of expert neurologists, genetic counselors, electrophysiologists, and data scientists.

### Translation of variant functional assessments using the FENICS ontology

For each experiment, we manually translated the reported results to the most precise FENICS terms possible. We assigned terms denoting abnormalities of a parameter only if the effect was reported as statistically significant (p<0.05) compared to WT. We included “normal” terms for a parameter only if the parameter was actually measured and not significantly different between the WT and mutant channels (p>0.05). We also assigned a single term denoting the experimenters’ conclusion as to the variant’s overall functional consequence as gain-of-function, loss-of-function, mixed/unclear, or normal. For example, an experiment on *SCN8A* p.M139I identified a significant left-shift in V_1/2_ of activation, but the slight increase in peak current density was within range of WT; accordingly, this experiment was assigned the terms “Moderate hyperpolarizing shift of voltage dependence of activation” (FENICS:0030), “Normal peak current” (FENICS:0096), and “Overall mixed function” (FENICS:0145).^34^

To include the full depth of functional information, we performed automated reasoning using the FENICS ontology as follows: for each abnormal measurement in an assessment, we assigned all higher-level, more general terms as well. For example, this process reflects the fact that a finding of “severe decrease in peak current” is also a finding of “decrease in peak current.” This process has been used previously, e.g., when conducting phenotype analysis using the HPO, to ensure that information from different sources and at different levels of specificity are harmonized for downstream analyses.^23,35-41^

### Association of functional consequences with variant class and location

The voltage-gated sodium channels have high sequence similarity, with most residues conserved across *SCN1A*, *SCN2A*, *SCN3A*, and *SCN8A*. Accordingly, each sequence position has been indexed to allow for comparison across these genes. For example, the arginine residues at position 1621 in *SCN3A* and position 1617 in *SCN8A* represent a conserved codon sequence. As such, these positions are mapped to the same index value.^42^ We labeled each variant with the index corresponding to its sequence position and defined subgroups of variants by position for subsequent analysis.

The voltage-gated sodium channels also have similar tertiary structures, consisting of four domains each containing six transmembrane segments, with the inactivation gate between domains III and IV. Within each domain, the segments serve similar roles, such as the voltage-sensing function of S4 and the pore-forming ion selectivity filter region between S5 and S6.^43,44^ We defined additional variant subgroups based on location within each domain or segment.

We then used the harmonized dataset of functional effects to compare frequencies of individual effects in specific subgroups to the remainder of the assessments. Using Fisher’s exact test with correction for multiple comparisons using False Discovery Rate (FDR) of 10%, we determined associations between FENICS-annotated functional changes and variants within individual segments and domains.

### Population-level analysis of variants in *SCN2A*

We compared the curated, functionally studied variants in *SCN2A* to our previously reported cohort of 413 individuals with *SCN2A*-related disorders and known disease-causing variants in *SCN2A*.^23^ We determined the ratio of those with missense variants whose variants were accounted for by existing functional data. We then assessed variants at conserved codons in *SCN1A/3A/8A*, as well as putative loss-of-function (pLOF) variants, defined as variants affecting splicing, nonsense variants, and frameshift variants. Using this expanded collection of variants, we estimated the proportion of individuals with *SCN2A*-related disorders for whom any variant functional information was available. This analysis was limited to *SCN2A,* as it is the only voltage-gated sodium channel gene with an established comprehensive patient cohort that includes every previously described individual, enabling a reliable proxy measure for the population prevalence of available functional information.

### Statistical analysis

All computations were performed using the R Statistical Framework. Statistical testing for associations is reported with correction for multiple comparisons using False Discovery Rate (FDR) of 10%. In cases where statistical significance was not reached after correction for multiple comparisons, findings remain on a descriptive level and are presented as odds ratios with 95% confidence intervals. Primary data for this analysis is available in the Supplementary material. Code for all analyses is available at github.com/helbig-lab/FENICS.

## Results

### Electrophysiological measurements in *SCN1A/2A/3A/8A* converge on 20 common parameters

In considering curation of electrophysiology data at scale, we reasoned that the functional consequence of a variant can be sufficiently described by precise, categorical information about well-defined biophysical parameters, such as peak current density. Based on consensus from an interdisciplinary panel of 18 researchers and clinicians, we identified 20 such commonly measured properties (**Table 1**). We recorded directional, nominally statistically significant (p < 0.05) differences in these parameters across 216 reports of 191 variants in *SCN1A* (n=74), *SCN2A* (n=66), *SCN3A* (n=18), and *SCN8A* (n=33), obtaining a baseline set of 1,484 annotations of features such as “hyperpolarizing shift in voltage dependence of activation” (**Table S1, Figure 1A-C**).

**Table 1.** Biophysical parameters.

|  |  |
| --- | --- |
| Peak current density ($I_{NaT}$) | Maximum current density (pA/pF) of current conducted by the channel |
| Rate of activation ($\tau_{act}$) | Time constant measuring how quickly or slowly the channel changes to the activated state with respect to time |
| Rate of deactivation ($\tau_{de}$) | Time constant measuring how quickly or slowly the channel closes upon repolarization with respect to time |
| Voltage dependence of activation ($V_{1/2act}$) | Half-maximal voltage, the potential at which 50% of channels are in the open state, reflecting the proportion of channels that are in activated states after being held at different membrane potentials/voltages |
| Slope factor of activation ($k_{act}$) | Slope factor indicating the steepness of voltage dependence of activation when plotted with respect to voltage after fitting to a Boltzmann function |
| Rate of fast inactivation ($\tau_{fi}$) | Time constant measuring how quickly or slowly the open channel changes to the inactivated state with respect to time |
| Rate of recovery from fast inactivation ($\tau_{rec,fi}$) | Time constant measuring how quickly or slowly the channel becomes available for activation after return to hyperpolarizing potentials to allow recovery from inactivation, i.e., the channel's inactivation gate reopens |
| Voltage dependence of fast inactivation ($V_{1/2fi}$) | Half-maximal voltage, the potential at which 50% of channels are in the fast inactivated state, reflecting the proportion of channels that are in fast inactivated states after being held at different membrane potentials/voltages |
| Slope factor of fast inactivation ($k_{fi}$) | Slope factor indicating the steepness of voltage dependence of inactivation when plotted with respect to voltage after fitting to a Boltzmann function |
| Persistent current ($I_{NaP}$) | Noninactivating current, the proportion of peak current that remains after the channel changes to the inactivated state |
| Gating-pore current | Presence of an alternative, abnormal pathway of ion flow created through the voltage-sensor domain ("omega current") resulting in abnormal depolarization due to additional cation flow, independent of the voltage-gated pore pathway |
| Channel modulation | Response of the channel to modulators, such as channel blockers or openers |
| Ion selectivity | Specificity of the channel for a single ion type |
| Resurgent current ($I_{NaR}$) | Current conducted due to the channel reopening in response to negative voltage changes after a normal transient current |
| Entry into slow inactivation ($\tau_{ent,si}$) | Time of onset of slow inactivation, i.e., entering an inactivated state following long (seconds) or high frequency depolarizations, reducing the proportion of channels available for opening |
| Rate of slow inactivation ($\tau_{si}$) | Time constant measuring how quickly or slowly the open channel changes to the slow inactivated state with respect to time |
| Rate of recovery from slow inactivation ($\tau_{rec,si}$) | Time constant measuring how quickly or slowly the channel becomes available for activation after return to hyperpolarizing potentials to allow recovery from inactivation, i.e. the channel's inactivation gate reopens |
| Voltage dependence of slow inactivation ($V_{1/2si}$) | Half-maximal voltage, the potential at which 50% of channels are in the slow inactivated state, reflecting the proportion of channels that are in slow inactivated states after being held at different membrane potentials/voltages |
| Slope factor of slow inactivation ($k_{si}$) | Slope factor indicating the steepness of voltage dependence of inactivation when plotted with respect to voltage after fitting to a Boltzmann function |
| Subthreshold current ($I_{NaS}$) | Continuous low-level conduction of ions by the channel |
| Ramp current ($I_{ramp}$) | Total amount of current measured during a steady depolarization over time |
| Use dependence (u.d.) | Attenuation of current conducted by the channel in response to repeated depolarization |

### Overall gain-of-function is heterogeneous, and loss-of-function homogeneous, in Na_V_ channels

To contextualize our curated functional results within the commonly used gain-of-function/loss-of-function paradigm, we first mapped the direction of the effect on each parameter based on its contribution to overall gain- or loss-of-function or overall normal function (**Figure 2A**). Among 182 experiments with measurable whole cell current, 74 (40.7%) had at least one defect contributing to gain-of-function and one contributing to loss-of-function, suggesting that resolution of these mutant channels as overall gain- or loss-of-function is not straightforward. However, only 36 of these experiments were reported by investigators as having mixed or unclear overall effect; an additional 18 assessments without bidirectional defects were considered mixed or unclear overall. Of the remaining assessments, 79 were reported overall gain-of-function and 72 overall loss-of-function.

We further explored these inferred functional effects by examining the most frequent changes from each variant type, representing effects that most commonly mediate the overall functional consequence. Among overall gain-of-function variants, the most frequent shifts were increased persistent current (f=0.52), hyperpolarizing shift of V_1/2_ of activation (f=0.46), and depolarizing shift of V_1/2_ of fast inactivation (f=0.42), with no single abnormality present in a commanding majority of assessments (**Figure 2B**). Similarly, the biophysical abnormalities with highest frequencies in the mixed/unclear category were hyperpolarizing shift of V_1/2_ of fast inactivation (f=0.43), slowing of recovery from fast inactivation (f=0.33), and increased persistent current (f=0.31, **Figure 2B**). In summary, variants with either gain-of-function and mixed or unclear overall effects have defects affecting multiple biophysical elements, such as persistent current and gating kinetics. No single biophysical abnormality drives functional changes in gain-of-function variants, which may represent an important insight for precision medicine approaches directed towards variants exhibiting gain-of-function mechanisms.

In contrast to gain-of-function variants, the most common features in loss-of-function variants were absence of current (f=0.47), decrease of peak current (f=0.40), and hyperpolarizing shift in V_1/2_ of fast inactivation (f=0.18, **Figure 2B**). Excluding variants where no cell current was measurable at all (n=34), most of these loss-of-function variants still involved a decrease in peak current density (f=0.76), indicating that peak current reduction is the predominant mechanism of loss-of-function in voltage-gated sodium channel variants.

### Electrophysiological studies on *SCN1A/2A/3A/8A* achieve a PS3_Strong ClinGen evidence level

Having identified the most common biophysical drivers of gain- and loss-of-function in available electrophysiology data, we sought to evaluate these parameters with respect to the PS3 criterion of the formal ACMG/AMP clinical variant interpretation guidelines.8 To this end, we applied a strategy consistent with the Bayesian framework recommended by the ClinGen Sequence Variant Interpretation (SVI) working group for modifying ACMG/AMP criteria.8,9,13,25

For each parameter, we initially computed a positive local likelihood ratio (LR^+^) as previously described^25^ for variant pathogenicity at every possible threshold value within our dataset. We analyzed data from 23 benign control variants in *SCN1A* and 63 pathogenic control variants (38 *SCN1A*, 6 *SCN2A*, 7 *SCN3A*, and 12 *SCN8A*, **Table S2**) for the four major drivers of gain- and loss-of-function: peak current density, voltage dependence of activation, voltage dependence of fast inactivation, and persistent current (**Figure 2**). For reduction in peak current density, reduction to less than 97.8% WT achieved *Supporting* evidence (LR^+^ = 2.22), less than 81.5% WT achieved *Moderate* evidence (LR^+^ = 5.00), and less than 74.2% WT achieved *Strong* evidence (LR^+^ = 20.0). Persistent current above 93% WT was considered *Supporting* evidence (LR^+^ = 2.10), while increases beyond 126% (LR^+^ = 5.25) and 135% (LR^+^ = 21.0) of WT achieved *Moderate* and *Strong* evidence, respectively. For voltage dependence of activation, the thresholds for *Supporting*, *Moderate*, and *Strong* were ±0.978 mV (LR^+^ = 2.10), ±2.15 mV (LR^+^ = 5.25), and ±2.20 mV (LR^+^ = 21. 0). Lastly, shifts in voltage dependence of fast inactivation of ±1.56 mV (LR^+^ = 2.22), ±2.96 mV (LR^+^ = 4.99), and ±4.10 mV (LR^+^ = 19.9) respectively represented *Supporting*, *Moderate*, and *Strong* thresholds (**Figure 3B-C**). As a result, voltage clamp results across parameters and evidence levels can be used within the PS3 ACMG/AMP criterion for the epilepsy sodium channels.

### *KCNQ2* voltage clamp studies reach PS3_Moderate and PS3_Strong ClinGen evidence levels

In order to extend our approach to a different type of voltage-gated ion channel with different experimental paradigms, we considered *KCNQ2*, which encodes the Kv7.2 protein and is the most common epilepsy-related potassium channel gene.^1,2,45^ In contrast to the sodium channels, KCNQ channels are tetrameric and non-inactivating. Available functional data comparing control population variants and pathogenic variants in *KCNQ2* included three important functional parameters: peak current density, voltage dependence of activation, and time constant of activation. Across measurements obtained under conditions mimicking the *KCNQ2* homozygous state of 80 disease-associated variants and 24 population variants, we determined that a peak current of less than 72.8% WT constitutes *Supporting* evidence (LR^+^ = 2.20), less than 61.6% WT *Moderate* (LR^+^ = 4.39), and less than 46.0% WT *Strong* (LR^+^ = 21.3, **Figure 4A-B**). Similarly, an increase in time constant of activation by more than 5% (LR^+^ = 2.31), 13% (LR^+^ = 5.01), and 24% (LR^+^ = 19.3) respectively represented *Supporting*, *Moderate*, and *Strong* evidence levels. For voltage dependence of activation, *Strong* evidence was not achieved, but the thresholds for *Supporting* and *Moderate* evidence were shifts of ±3.59 mV (LR^+^ = 2.11) and ±5.77 mV (LR^+^ = 4.50), respectively (**Figure 4B**).

As expected, thresholds for the heterozygous state were closer to WT than those for the homozygous state. Specifically, we identified *Supporting*, *Moderate*, and *Strong* evidence cutoffs of 91.0% (LR^+^ = 2.20), 81.5% (LR^+^ = 5.40), and 68.4% (LR^+^ = 19.2) WT for peak current density. Increase in time constant of activation by more than 5% (LR^+^ = 2.23) and 9% (LR^+^ = 4.45) of WT, and shift in voltage dependence of activation of more than ±3.59 mV (LR^+^ = 2.25) and ±5.77 mV (LR^+^ = 5.67), achieved *Supporting* and *Moderate* evidence respectively; neither of these parameters achieved a level of *Strong* with available data (**Figure 4C**).

To maintain consistent granularity across parameters and channels, for parameters that achieved only up to *Moderate* evidence, we approximated a cutoff for the most severe shifts in these parameters based on the distribution of benign variants. Specifically, across all parameters for which *Supporting*, *Moderate*, and *Strong* thresholds could be calculated, these thresholds mirrored the 50^th^, 80^th^, and 95^th^ percentile of benign variant measurements (**Figure 4C**). Accordingly, we used the same 95^th^ percentile to estimate cutoffs for a severe shift in voltage dependence of activation at ±3.24 mV in the heterozygous state (LR^+^ = 16.5) and at ±7.58 mV in the homozygous state (LR^+^ = 17.0), as well as severe slowing of activation at a time constant of more than 13% in the heterozygous state (LR^+^ = 17.1, **Figure 4B-C**). In summary, evaluation of *KCNQ2* electrophysiology represents an extension of our framework beyond the sodium channels to allow assessment of more complex experiments.

### The biomedical ontology for curation of electrophysiological data contains 152 concepts

Although we had calibrated several biophysical parameters for use with ACMG/AMP criteria, a critical gap in directly applying these findings in the clinical setting lies in the practical difficulty of interpreting heterogeneous reporting of functional data in the literature. Accordingly, we compiled our rich categorical classifications of biophysical parameters and threshold values into a biomedical ontology, the Functional Electrophysiology Nomenclature for Ion Channels (FENICS, available at bioportal.bioontology.org/ontologies/FENICS). This represents a set of 152 hierarchical, standardized labels to capture the functional consequence of a variant, with strictly defined terminology that, for the calibrated parameters, can map directly to ACMG/AMP evidence levels (**Figure 5A**).

Based on expert consensus, 20 parameters, including persistent current, voltage dependence of activation, and peak current density, are sufficiently distinct and elementary to permit a hierarchical classification (**Table 1**). However, some features reported in experimental results represent more complex phenomena that are not easily decomposed into discrete biophysical changes. This necessitated a second, structurally similar sub-ontology for terms such as ramp current and use dependence.

Additionally, we accounted for similar effects of different biophysical parameters. Intuitively, a decrease in peak current and a slowing of activation are conceptually related, as both changes reflect reduction in channel function. In particular, the slowing of activation contributes to a loss of function of the Na_V_1.2 channel because the channel does not open for longer durations, allowing less sodium to cross the plasma membrane. Accordingly, our dictionary connects each electrophysiological change implicitly to a term indicating whether it contributes to a gain-of-function or loss-of-function effect in overall ion channel function (**Figure 5B**).

Lastly, investigators typically infer the overall functional consequence of a variant from its electrophysiological data. However, as we found above, experiments frequently show changes in both gain- and loss-of-function directions, even when investigators conclude that a variant results in overall gain- or loss-of-function. For example, for the *SCN8A* p.R1872W variant, the combination of mild slowing of fast inactivation, mild hyperpolarizing shift in voltage dependence of activation, and moderate increase in slope of activation is assumed to result in an overall gain-of-function.^46^ Thus, we considered the investigators’ conclusion to be a distinct concept from individual biophysical changes and included them in a separate subontology (**Figure 5B**).

### Curation of 216 assessments yields 4,272 functional annotations across *SCN1A/2A/3A/8A*

We retrieved 216 total functional assessments in the literature for missense variants in *SCN1A* (n=82), *SCN2A* (n=72), *SCN3A* (n=18), and *SCN8A* (n=44, **Figure 1A-B**). For the 216 functional assessments, we translated a total of 1,484 FENICS terms with a range of 1-15 terms and median of 8 terms per experiment (**Table S1**). In total, we used 89 unique translated terms of a possible 152, or 58.6% of terms within the entire dictionary. Of these assigned terms, 781 of 1,484 denoted abnormalities, including 70 unique terms with a median 3 terms and range of 1-13 per experiment. The other 703 of 1,484 translated terms referred to normal measurements, or measurements within statistical range of WT controls in a given experiment. We used a total of 19 unique normal terms with a median of 4 terms and range of 1-12 per experiment. The most commonly assigned terms were normal terms, including “Normal peak current” (FENICS:0096, f=0.47), “Normal slope of activation” (FENICS:0036, f=0.44), and “Normal slope of fast inactivation” (FENICS:0074, f=0.42). The most common abnormal terms were “Absence of peak current” (FENICS:0083, f=0.16) and “Severe increase of persistent current” (FENICS:0043, f=0.15).

Following automated inclusion of inferred terms, we obtained a harmonized dataset reflecting the complete collection of functional alterations across these curated assessments, which included 4,272 total calculated terms. Of the 152 possible distinct terms, we used 123 (80.92%) in this curation. There was a median of 20 terms and a range of 1-59 terms per experiment. Within this more comprehensive collection, among the most common functional categories were “Effect on fast inactivation” (FENICS:0037, f=0.66), “Normal peak current” (FENICS:0082, f=0.47), and “Effect on peak current” (FENICS:0020, f=0.44), reflecting that these features are the most common mechanisms driving the overall functional effect of variants.

Within the curated set of 1,484 annotations, there are 330 terms across 169 variants (88.5%) that are directly usable at *Supporting*, *Moderate*, or *Strong* levels for variant classification. In the FENICS ontology, terms denoting mild, moderate, and severe changes to certain parameters map to these modified ACMG/AMP thresholds. For instance, a mild hyperpolarizing shift in voltage dependence of activation (FENICS:0029) is defined as a left shift of between 0.978 mV and 2.15 mV, which corresponds to *Supporting* evidence in clinical variant interpretation; and a severe decrease in peak current (FENICS:0087) is defined as a reduction beyond 74.2% of WT current density and thus indicates *Strong* evidence of variant pathogenicity. There were 22 variants for which no available assessment included our calibrated parameters for the PS3 criterion, including nine which were considered to be overall normal with respect to voltage clamp. Among the main abnormalities in the remaining 13 assessments were slope factor of fast inactivation (f=0.31), rate of fast inactivation (f=0.31), and rate of recovery from fast inactivation (f=0.23). Given the low frequency of these abnormalities, we were underpowered to calibrate these parameters per the ACMG/AMP framework. As more, consistent measurements of these parameters become available in future experiments, we expect more rigorously defined thresholds for severity to be achievable across the full breadth of electrophysiological study of voltage-gated sodium channel variants.

Our final dataset of 1,484 annotations is accessible in the functional evidence section of the ClinVar database for a given variant. As a result, FENICS annotations can be used within the ACMG/AMP PS3 criterion for an epilepsy ion channel variant in an accessible, standardized, digital format.

### FENICS captures known genotype-function associations in voltage-gated sodium channels

Given the known structure-function relationships and topological similarity across these ion channels,^42,43^ we applied FENICS to map functional alterations to common channel structural elements at the levels of transmembrane repeat domain and individual transmembrane segment. Following Fisher’s exact test with correction for FDR of 10%, we found significant associations between missense variants in S1 and hyperpolarizing shifts of voltage dependence of activation (FENICS:0027, p=0.0019, OR=17.4, 95%CI=2.04-813); S4 and increase in slope factor or fast inactivation (FENICS:0073, p=0.0058, OR=3.81, 95%CI=1.34-10.3); S5 and depolarizing shifts of voltage dependence of slow inactivation (FENICS:0116, p=0.0040, OR=27.8, 95%CI=2.12-1503); S5-6 and overall loss-of-function (FENICS:0141, p=0.0012, OR=7.50, 95%CI=1.59-71.9) and absent current (FENICS:0083, p=0.0001, OR=10.7, 95%CI=2.84-44.9; and S6 and absent current (FENICS:0083, p=0.0060, OR=3.55, 95%CI=1.31-9.21; **Figure 6A, Table S3**). The associations with rate and slope of inactivation in particular reflect known functions of segment 4 in voltage sensing.^43^ Similarly, the absence of current in S5-6 and S6 variants is explained by the involvement of these regions in pore formation.^43^ These findings also parallel prior work showing clustering of loss-of-function variants at the pore and gain-of-function mechanisms at the voltage sensing domain.^47,48^

**Figure 6.**
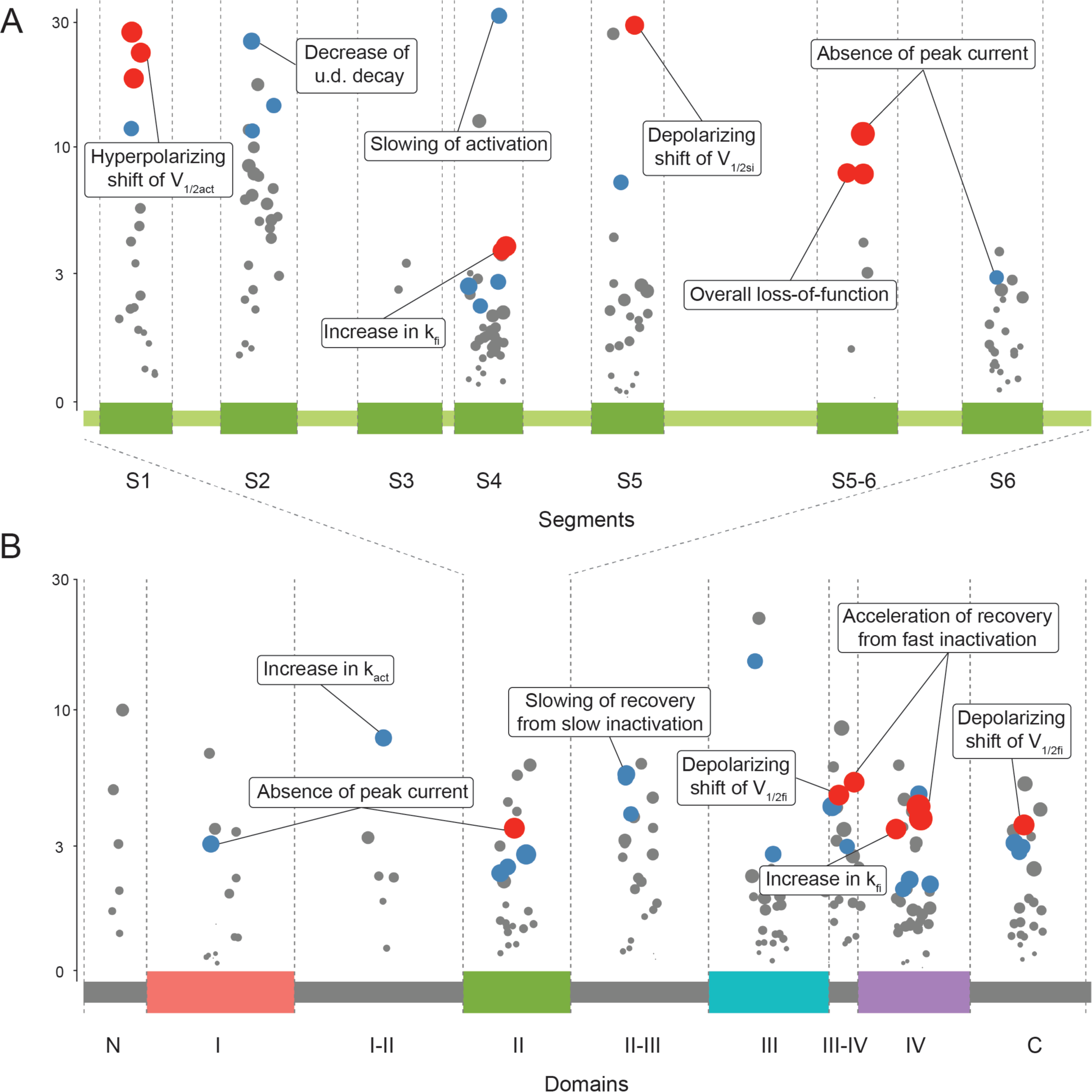
Association between specific biophysical changes and common structural features of voltage-gated sodium channels, i.e., transmembrane segments (A) and repeat domains (B). Points are plotted along the protein primary sequence. Point size indicates p-value and height indicates odds ratio. Red points are associations significant after correction for FDR of 10%; blue points are associations nominally significant with p < 0.05; gray points are non-significant associations with p ≥ 0.05. V1/2act = voltage dependence of activation, u.d. = use dependence, kfi = slope factor of fast inactivation, V1/2si = voltage dependence of slow inactivation, kact = slope factor of activation, V1/2fi = voltage dependence of fast inactivation.

Among transmembrane repeat domains, variants in domain II were significantly associated with absent current (FENICS:0083, p=0.0030, OR=3.53, 95%CI=1.44-8.45); the domain III-IV linker with depolarizing shifts of voltage dependence of fast inactivation (FENICS:0064, p=0.0001, OR=14.5, 95%CI=3.43-61.2) and faster recovery from fast inactivation (FENICS:0053, p=0.0036, OR=5.29, 95%CI=1.55-17.2); domain IV with faster recovery from fast inactivation (FENICS:0053, p=0.0005, OR=4.27, 95%CI=1.78-10.4) and increased slope factor of fast inactivation (FENICS:0073, p=0.0039, OR=3.50, 95%CI=1.38-8.90); and the C-terminal region with depolarizing shifts of voltage dependence of fast inactivation (FENICS:0060, p=0.0011, OR=3.63, 95%CI=1.46-8.93; **Figure 6B, Table S3**). Here, altered inactivation parameters are as would be expected of the domain III-IV linker, given its role in inactivation gating.^43^ In summary, the distributions of functional annotations across the channel provide validation of FENICS in sufficiently capturing structural-functional relationships from established functional domains and motifs in sodium channels and suggest potential value for this ontology in categorical assessment of channel defects at larger scales.

### Functional consequences can apply to a majority of individuals with *SCN2A*-related disorders

Functional data are not just applicable in the diagnostic space through the ACMG/AMP PS3 criterion, but also are increasingly useful in therapeutic decision-making. Accordingly, to assess the total potential impact of already available electrophysiological data, we examined the curated dataset at a disease population level by projecting the results of 72 experiments on 66 *SCN2A* variants onto a cohort of 413 individuals with *SCN2A*-related disorders. Excluding protein-truncating variants and large deletions, which are presumed to result in complete loss-of-function, we found that 124/343 (36.2%) of individuals had missense variants with existing functional evidence. In particular, 103/343 (30%) of individuals had one of 19 recurrent missense variants, including 10 functionally studied variants accounting for 21% of individuals (**Figure 7**).

**Figure 7.**
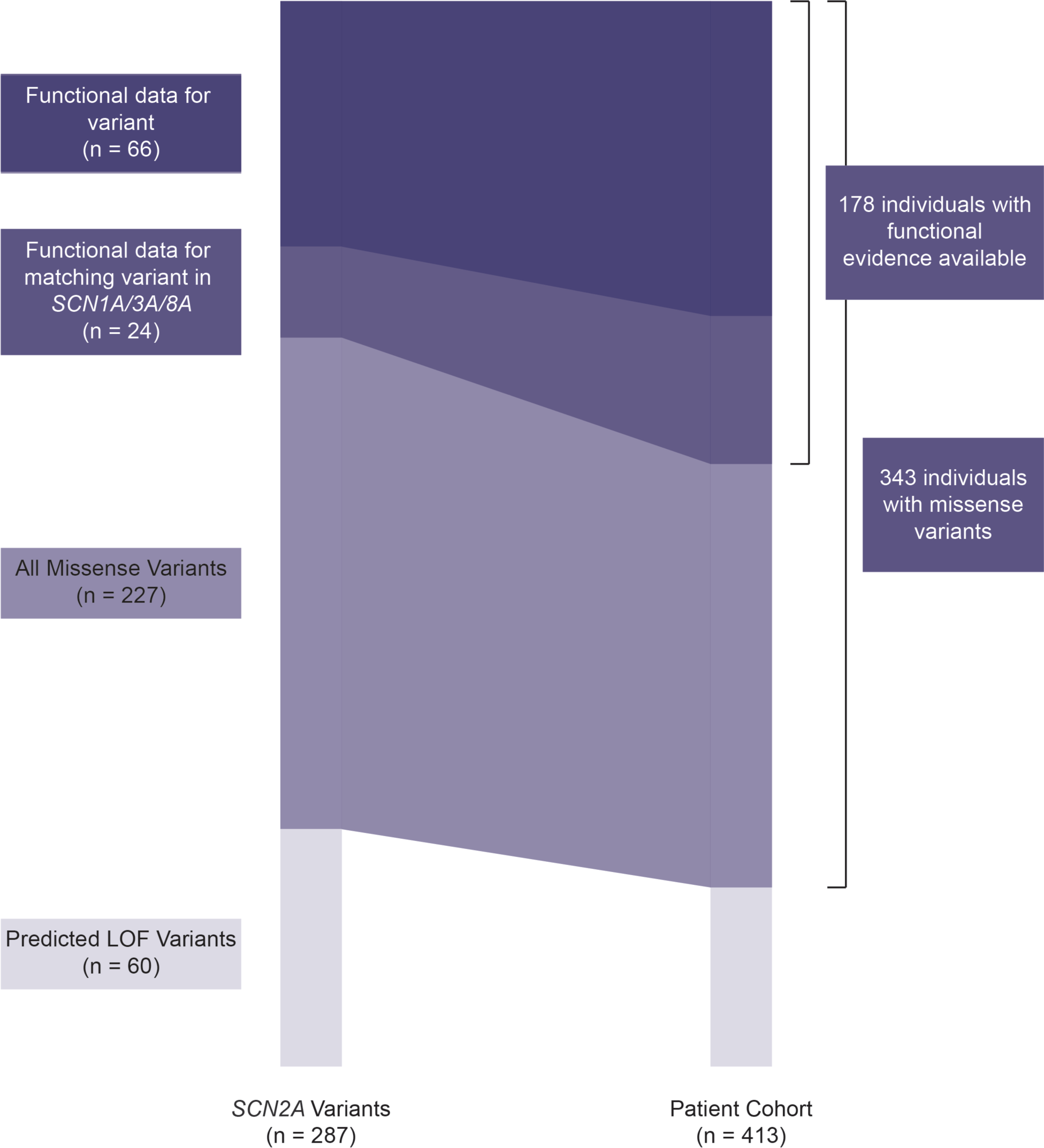
Projection of FENICS-curated *SCN2A* variants (n=66) to 413 individuals reported with *SCN2A*-related disorders. More than 1 in 3 individuals with missense variants have functional data available for their variant, while more than half have functional data available for at least one variant affecting the corresponding amino acid across *SCN1A/2A/3A/8A*.

Next, we included disease-causing variants at conserved codons across the four channels *SCN1A/2A/3A/8A*. This accounted for 24 additional variants and 54 individuals with some related functional information. Including the entire disease cohort, variants with existing functional information across the Na_V_ channels accounted for 52% of individuals with missense variants causing *SCN2A*-related disorders (**Figure 7**). By establishing a population-level understanding of the scope of functional data in *SCN2A*, this dispels the notion that functional information is unknown for most individuals, which is critical as functional data on missense variants can contribute substantially to both diagnosis of *SCN2A*-related disorders and the clinical actionability of such a diagnosis.

## Discussion

Here, using the Bayesian approach adopted by the ClinGen Sequence Variant Interpretation Working Group,^9,13,25^ we determined the levels at which electrophysiological data can be used for clinical variant interpretation of several voltage-gated ion channel genes in neurodevelopmental disorders. We compiled functional data into the newly built FENICS ontology and show that its 152 consensus terms are sufficiently precise to describe functional impact for use in the ClinGen PS3 framework.

Disease-causing variants in genes encoding voltage-gated ion channels such as *SCN1A*, *SCN2A*, *SCN3A*, *SCN8A*, and *KCNQ2* are the most common causes of genetic epilepsies.1,2 Voltage clamp studies on these channels have been performed for decades and remain a standard assay; thus, they are continuously and increasingly a part of the clinical decision-making space for these genetic conditions.21,33 Yet, variable and nonstandard reporting and cataloguing has led to limited analysis of the entirety of available data and represents a major impediment to accessibility of these results in the clinic.

In our work, we reasoned that a data harmonization approach to electrophysiological data would facilitate deeper discoveries, as evidenced by the analogous impact of the Human Phenotype Ontology (HPO) in genotype-phenotype studies.30,49,50 Since its first release, the HPO has been used by us and others to comprehensively evaluate clinical presentations, identify novel disease genes, and carry out extensive longitudinal phenotyping efforts.23,35-41

To surmount the barrier to incorporating functional data in precision care of individuals with channelopathies, we developed the FENICS ontology to act as a bridge across electrophysiology labs, ClinVar, ClinGen, and the clinic in voltage-gated channelopathies. Combining standardized parameters from FENICS with quantitative electrophysiological data, we determined threshold values for *Strong*, *Moderate*, and *Supporting* weights for decrease in peak current, increase in persistent current, and shifts in voltage dependence of activation or inactivation, as per the ClinGen SVI Working Group criteria.^8,9,13^ Of note, supporting thresholds tended to be well within the variability of wildtype channels, suggesting more caution may be warranted in applying a PS3_Supporting criterion from our calibration than in higher levels of evidence. Analogous to the HPO’s applicability across disease domains, we calibrated PS3 in a different epilepsy-related channel gene, *KCNQ2*, with distinct parameters and thresholds, illustrating the utility of FENICS across ion channels. However, the potential direct benefit of all these raw threshold values may be limited, as finding and interpreting heterogeneously presented data from research laboratories can be difficult in the context of clinical genetic testing. Accordingly, the weighted thresholds are also mapped to existing FENICS terms at levels of “mild,” “moderate,” and “severe” changes to a given parameter. Beginning March 2022, FENICS is an accepted format for describing functional conseqences of variants in ClinVar, and 271 variants with FENICS terms have been deposited to date. Therefore, FEINICS provides a standardized and publicly available framework for simplified variant interpretation that provides accessible precise functional information to clinical providers.

One feature of our work was the harmonization of electrophysiological results across not only several distinct reports, but also different sodium channel genes, fueled by the high sequence similarity between the four neuronally expressed voltage-gated sodium channels. An emerging body of work highlights how inference across these channels can accurately represent broad functional changes.^42,48^ In a specific example, this approach has already helped define the clinical-genetic spectrum of p.R1636Q, a recurrent reported gain-of-function *SCN1A* variant.^21,22^ Our FENICS dataset recapitulated several structure-function relationships among sodium channels, demonstrating that the ontology captures functional consequences granularly enough to reflect some underlying biology. Moreover, as a result of our cross-channel approach, we were able to apply our calibration beyond *SCN1A*, which accounts for our benign variant dataset, to an additional 117 variants in *SCN2A/3A/8A*, which are the causative genes in up to an additional 11% of all genetic epilepsies in large exome studies.^1,2^ Given the applicability of some functional insight for a majority of reported individuals with disease-causing *SCN2A* variants, as well as the rise of automated electrophysiological approaches in the research space,^15-18^ we expect that systematic functional evaluations will facilitate diagnosis and expand clinical trial readiness.

The 191 experimental sodium channel variant assessments harmonized in our study account for a significant proportion of patients seen in a clinical setting. For example, we estimate that clinical care of nearly half of all individuals with early-onset *SCN2A*-related disorders can be informed by existing functional data on their variant alone. Furthermore, given that variants at identical sites across brain-expressed voltage-gated sodium channels are largely functionally consistent, an even larger proportion of diagnosed individuals already have some functional information to guide clinical care and therapeutic decision-making.

Humans possess 79 genes encoding potassium channel pore-forming subunits, making them the most diverse ion channel gene superfamily,51 and at least 19 of these genes have been linked to epilepsy and/or NDD to date.52 To begin to test the broader potential utility of our approach for evaluation of potassium channel genes, we applied it to *KCNQ2,* using a recent dataset where 80 clinical and 24 control population variants had been studied by patch-clamp in parallel under standardized conditions.

In summary, our data demonstrates the utility of curating electrophysiological studies in voltage-gated ion channels to satisfy criteria for formal clinical variant interpretation. There remains a pressing need to generate electrophysiological data on novel, unstudied variants, but our findings emphasize that there is also a significant amount of data that could be incorporated into clinical management of individuals with these disorders. By establishing a systematic and standardized language for existing functional data, we expand the availability and promote the clinical actionability of ion channel electrophysiology.

## Acknowledgements

I.H. was supported by NINDS (K02NS112600, U24NS120854-01, U54NS108874, R01NS131512, R01NS127830), The Hartwell Foundation (Individual Biomedical Research Award), Simons Foundation Autism Research Initiative, the Eunice Kennedy Shriver National Institute of Child Health and Human Development through the Children’s Hospital of Philadelphia and the University of Pennsylvania (U54HD086984), the German Research Foundation (HE5415/3-1, HE5415/5-1, HE5415/6-1, Research Unit FOR-2715: HE5415/7-2, HE5415/7-1), the National Center for Advancing Translational Sciences of the NIH (UL1TR001878), the Dravet Syndrome Foundation Genetic RFA, the Institute for Translational Medicine and Therapeutics (ITMAT) at the Perelman School of Medicine of the University of Pennsylvania, and by Children’s Hospital of Philadelphia through the Epilepsy NeuroGenetics Initiative (ENGIN). D.L.S. was supported in part by a research grant from Science Foundation Ireland (SFI) under Grant Number 21/RC/10294_P2 and co-funded under the European Regional Development Fund and by FutureNeuro industry partners. The contribution of S.L., C.B., U.H. and H.L. was supported by the German Research Foundation (Research Unit FOR-2715: LE1030/15-2, LE1030/16-2, HE8155/1-2) and the Federal Ministry for Education and Research (BMBF, consortium on rare neurological ion channel disorders Treat-ION, 01GM2210A). R.K. was supported by the Fonds National de la Recherche Luxembourg (FNR; Research Unit FOR-2715, FNR grant INTER/DFG/21/16394868 MechEPI2). E.C.C. was supported by the Miles Family Fund, the Jack Pribaz Foundation, and U54NS108874-1. The work of M.J.L. and J.H. was supported by the National Center for Biotechnology Information of the National Library of Medicine (NLM), National Institutes of Health.

## Web Resources

Online Mendelian Inheritance in Man, http://www.omim.org

Human Phenotype Ontology, https://hpo.jax.org

Functional Electrophysiology Nomenclature for Ion Channels, https://bioportal.bioontology.org/ontologies/FENICS

## Data and Code Availability

The published article includes all datasets generated during this study. The code generated during this study is publicly available at github.com/helbig-lab/FENICS.

